# Understanding the limit of open search in the identification of peptides with post-translational modifications — A simulation-based study

**DOI:** 10.1101/289710

**Authors:** Jiaan Dai, Fengchao Yu, Ning Li, Weichuan Yu

**Author notes:** These authors contributed equally to this work (Jiaan Dai, Fengchao Yu).

## Abstract

**Motivation:** Analyzing tandem mass spectrometry data to recognize peptides in a sample is the fundamental task in computational proteomics. Traditional peptide identification algorithms perform well when identifying unmodified peptides. However, when peptides have post-translational modifications (PTMs), these methods cannot provide satisfactory results. Recently, Chick *et al.*, 2015 and Yu *et al.*, 2016 proposed the spectrum-based and tag-based open search methods, respectively, to identify peptides with PTMs. While the performance of these two methods is promising, the identification results vary greatly with respect to the quality of tandem mass spectra and the number of PTMs in peptides. This motivates us to systematically study the relationship between the performance of open search methods and quality parameters of tandem mass spectrum data, as well as the number of PTMs in peptides.

**Results:** Through large-scale simulations, we obtain the performance trend when simulated tandem mass spectra are of different quality. We propose an analytical model to describe the relationship between the probability of obtaining correct identifications and the spectrum quality as well as the number of PTMs. Based on the analytical model, we can quantitatively describe the necessary condition to effectively apply open search methods.

**Availability:** Source codes of the simulation are available at http://bioinformatics.ust.hk/PST.html.

**Supplementary information:** Supplementary data are available at *Bioinformatics* online.

## 1 Introduction

Mass spectrometry (MS) is currently the most important and versatile tool in large-scale proteomics (Aebersold and Goodlett, 2001; Yates *et al.*, 2009). During the past two decades, more and more protein analytical strategies have used mass spectrometry in the analysis of complex protein samples (Aebersold and Mann, 2003; Domon and Aebersold, 2006). To determine the amino acid sequences of peptides, tandem mass spectrometry is usually used as the key technique for peptide identification (Han *et al.*, 2008). Correspondingly, a large number of methods for peptide identification have been developed.

Database search is a type of method for peptide identification (Eng *et al.*, 1994, 2013; Cottrell and London, 1999; Tanner *et al.*, 2005; Wang *et al.*, 2007; Kim and Pevzner, 2014; McIlwain *et al.*, 2014). It uses a sequence database as a reference, and labels the second stage mass spectra (MS2) with known sequences based on the similarity scores between the query spectra and the sequences in the database. Database search provides accurate identification, but can only identify peptides that are already in the database.

*De novo* sequencing is another type of method (Dancik *et al.*, 1999; Chen *et al.*, 2001; Frank and Pevzner, 2005). It builds a spectrum graph from the MS2 spectrum, and infers the amino acid sequence by traversing the graph. *De novo* sequencing is able to identify novel sequences, but cannot provide results as accurately as database search.

Spectral library search is a third type of method (Lam *et al.*, 2007, 2008; Ahrné *et al.*, 2011). It uses a spectral library as a reference, and identifies experimental MS2 spectra based on the similarity scores between the query spectra and the library spectra. Spectral library search captures common patterns of peptides, but needs to store identified peptides in the spectral library.

Database search is widely used in peptide identification for its efficiency and robustness. When identifying peptides with post-translational modifications (PTMs), a number of computational methods have been proposed, on top of the database search framework. Tsur *et al.*, 2005 proposed the MS-Alignment algorithm which uses a dynamic programming (DP) approach, allowing jumps across an unmatched amino acid to conduct an error-tolerant spectral alignment between a sequence and an MS2 spectrum. Na *et al.*, 2012 proposed a different DP-based alignment algorithm using peptide sequence tags, which have been widely used in the proteomics community (Mann and Wilm, 1994; Tabb *et al.*, 2003; Frank *et al.*, 2005; Tanner *et al.*, 2005; Cao and Nesvizhskii, 2008; Kim *et al.*, 2009a,b), to measure the similarity between a sequence and an MS2 spectrum. Besides DP-based alignment algorithms, other proposed methods usually assume that only one unknown PTM will exist (Hansen *et al.*, 2005; Chalkley *et al.*, 2008; Chen *et al.*, 2009). Then, the mass shift of this unknown PTM can be inferred by subtracting the mass of the candidate peptide from the observed precursor mass.

However, there are some remaining issues that have not yet been fully addressed: (1) The number of PTMs that can be handled is still limited; (2) the accuracy of the identification results is not as good as the algorithms that enumerates modified variants.

To address these issues, Chick *et al.*, 2015 proposed an open search method for the identification of peptides with PTMs. The theoretical spectra without PTMs are directly compared with the query spectra using the XCorr scoring function (Eng *et al.*, 1994, 2008) in a commonly used database search procedure (Eng *et al.*, 1994). To include more modified candidates, they set the MS1 tolerance at 500 Da. MSFragger (Kong *et al.*, 2017) further accelerated the spectrum-based open search by using a novel fragment-ion indexing method. Recently, Yu *et al.*, 2016 proposed another open search approach named PIPI, which calculates the similarity using transformed tag vectors. The transformed tag vectors are extracted from the query spectra and the database sequences. PIPI uses the tag-based open search in its first stage identification, and finally identifies more peptide-spectral matches (PSMs) with less computing time compared with existing methods, e.g., MODa (Na *et al.*, 2012).

While Chick *et al.*, 2015 and Yu *et al.*, 2016 empirically showed promising results on several datasets, the feasible regions where the spectrum-based and tag-based open search works still remain unclear. When open search provides different interpretations compared with existing methods, we don’t know how reliable these interpretations are, although the false discovery rates (FDRs) of different interpretations have already been controlled at the specified level. Therefore, it is necessary to figure out when the open search can provide accurate identification.

In this paper, we carry out a simulation-based analysis on open search methods. To facilitate the quantitative analysis with known ground truth, we use simulated datasets with explicit assumptions. As real data will suffer from inappropriate dissociation energy, impurities, and experimental variations, we propose models of these common effects in the simulation. By conducting large-scale simulations, we obtain the performance trend of both spectrum-based and tag-based open search methods when the MS2 spectra are of different qualities. We further provide an analytical model to describe the relationships between the probability of obtaining correct identifications and the spectrum quality as well as the PTM condition. The evaluation of the proposed model shows that it can accurately describe the performance trend based on the given parameters. From the model, we are able to obtain the necessary condition to effectively apply both types of open search methods in peptide identification.

The rest of the paper is organized as follows. We first introduce the essential procedures of the spectrum-based and tag-based open search methods. Then, we explain the procedure to conduct simulations and models of common effects when generating simulated datasets. After that, we present the simulation results, and the analytical model to describe the performance trend of the spectrum-based and tag-based open search. We then evaluate the analytical model using independent testing datasets. Finally, we conclude the paper with some discussions.

## 2 Method

### 2.1 Open search for PTM identification

The basic idea of open search is to enlarge the tolerance of the precursor mass (MS1 tolerance) to include potentially modified candidate sequences but without considering PTMs when generating the theoretical representations of database sequences during the search (Chick *et al.*, 2015; Yu *et al.*, 2016; Kong *et al.*, 2017). Typically, the enlarged MS1 tolerance is set at *±*500 Da (Chick *et al.*, 2015) or *±*250 Da (Yu *et al.*, 2016).

In brief, open search methods can be classified into two categories: (1) spectrum-based approach (Chick *et al.*, 2015; Kong *et al.*, 2017) and (2) tag-based approach (Yu *et al.*, 2016). The difference between these two categories is the way to compute the similarity between a query spectrum and the database sequence. The spectrum-based approach generates the original theoretical spectrum of the database sequence without considering any PTMs, and then computes the similarity using the specified scoring function (e.g., XCorr (Eng *et al.*, 1994, 2008)) on top of the spectrum vectors. The tag-based approach generates the tag vectors of the database sequence and the tag vectors of query spectra by applying tag extraction algorithms, and then computes the similarity using the specified scoring function (e.g., cosine similarity (Yu *et al.*, 2016)) on top of the tag vectors.

### 2.2 Simulation approach

#### 2.2.1 Workflow

Instead of using real datasets in the analysis, using simulated datasets allows us to have detailed control of the experimental conditions and spectrum qualities. Even though we have the information about the general conditions in real experiments, the individual parameters of each spectrum are out of our control. For example, it is difficult to know whether a certain peptide is perfectly fragmented or not during the collision-induced dissociation. But in simulations, we are able to know the ground truth about how well a certain simulated peptide is fragmented. Moreover, we can cover more possible conditions using simulation.

To simplify the procedure and remove the effects introduced by scoring functions using intensity information, we use the binarized form of the spectrum. Therefore, in both spectrum-based and tag-based open search, the similarities are normalized dot products (a.k.a. cosine similarity) of binary vectors, i.e., setting the value to one if the corresponding m/z peak or tag exists. In addition, we only consider *b* and *y* ions with charge 1, and assume all precursors have one charge in the simulations. We conduct the simulation as follows:

1. Generate the number of PTMs from Poisson distribution (details in Section 2.2.2).
2. Uniformly draw a peptide from the database with a length larger than or equal to the number of PTMs as the base peptide of the simulated spectrum.
3. Generate random mass shifts to obtain the modified theoretical spectrum. The model for generating random mass shifts is stated in Section 2.2.2.
4. Generate noise peaks in the spectrum. The model for noise peak generation is stated in Section 2.2.3.
5. Delete some true peaks to mimic the effect of missing peaks in the spectrum. The model for true peak deletion is stated in Section 2.2.4. If some noise peaks are within the tolerance range of the true peak to be deleted, we delete these noise peaks at the same time. By doing so, we guarantee that we will not observe any peaks in the anticipated m/z position of this true peak. After this step, we obtain a simulated spectrum.
6. Search the simulated spectrum as a normal spectrum against the database (with a decoy database) using the spectrum-based and tag-based open search.
7. Collect the search results and check the consistence between the reported identification and the original base peptide. PTM localization is a follow-up task after the open search. Thus, we only examine the backbone peptide sequence here.
8. Go back to the first step to start the simulation of the next spectrum.

We need several models to control the behaviors of random mass shifts (mimicking the effect of PTMs), the behaviors of noise peaks (mimicking the effect of uninterpreted peaks in the identified spectrum), and the behaviors of missing peaks (mimicking the effect of unmatched theoretical ions). Below we introduce these models in detail.

#### 2.2.2 Model of mass shifts

We generate random mass shifts within the MS1 tolerance range to simulate the effect introduced by unknown PTMs. Given the number of mass shifts, we uniformly choose the modified amino acids from the base peptide without replacement. For a modified amino acid with mass *M*_*a*_, we generate a mass shift value from *U* (*−M*_*a*_, *ϵ*_1_), where ϵ_1_ is the MS1 tolerance. The precursor mass of the modified spectrum should be within the *±E*_1_ range of the mass of the base peptide; otherwise, the simulation tool will re-draw new realizations about the mass shifts. We assume that the number of random mass shifts follows a Poisson distribution with parameter *λ*, where *λ* describes the average number of random mass shifts in the simulated spectra.

#### 2.2.3 Model of uninterpreted peaks

The ratio of the number of noise ions (i.e., uninterpreted peaks in the spectrum) to the number of theoretical ions is denoted as *r*_*n*_. Given a parameter *r*_*n*_ and a theoretical spectrum, the number of noise peaks in the simulated spectrum is ⌊*nr*_*n*_⌋, where *n* denotes the number of theoretical ions. ⌊*nr*_*n*_⌋ is the largest integer smaller than or equal to *nr*_*n*_. The positions of noise peaks in the *m/z* axis follows a uniform distribution *U* (0, *P M*), where *P M* is the precursor mass of the simulated spectrum.

#### 2.2.4 Model of missing peaks

The ratio of the retained theoretical ions is denoted as *r*_*s*_. The proportion of missing peaks due to incomplete fragmentation, different ionization efficiencies, and different ionization potentials then equals 1 *- r*_*s*_. Given a parameter *r*_*s*_ and a theoretical spectrum, the number of missing peaks is ⌊(1 *- r*_*s*_)*n*⌋, where *n* is the number of theoretical ions, Every theoretical ion has the same probability of missing.

## 3 Results

### 3.1 Simulation results

To generate a simulated spectrum, we need to specify three parameters *r*_*s*_, *r*_*n*_ and *λ*. For each parameter combination (*r*_*s*_, *r*_*n*_, *λ*), we generate *N* = 40000 simulated spectra, and use the spectrum-based and tag-based open search methods to analyze the simulated spectra. We use the target-decoy approach with shuffling sequences (Elias and Gygi, 2007) and control the FDR at 0.05 at the PSM level for each parameter combination. Only when a PSM passes the FDR threshold will it be regarded as an identification. We scan the parameter space from *r*_*s*_ = 0 to *r*_*s*_ = 1 (sampling 11 points), from *r*_*n*_ = 0.1 to *r*_*n*_ = 100 (sampling 16 points), and from *λ* = 0.0 to *λ* = 3.75 (sampling 16 points), to investigate the performance when the MS2 spectra are of different qualities. When *λ* = 0, we fix the number of random mass shifts to be zero. We use the *Homo sapiens* protein database (downloaded on 2016.03.10 from UniProtKB/SwissProt, 20198 proteins) in these simulations. The database is digested with the trypsin cleavage rule. No missed cleavages are allowed. The allowed range of the peptide mass is [700, 5000] Da. The MS1 tolerance is 250 Da, and the MS2 bin size is 0.02 Da. We set the tag length to be equal to 3, which was empirically decided by Yu *et al.*, 2016. We collect the empirical probabilities of obtaining correct identifications in different parameter combinations, where the empirical probability of obtaining correct identifications is calculated by #{Correctly Identified Spectra} / #{Total Spectra}. The simulation results are shown in Fig. 1 and Fig. 2.

**Fig. 1:**
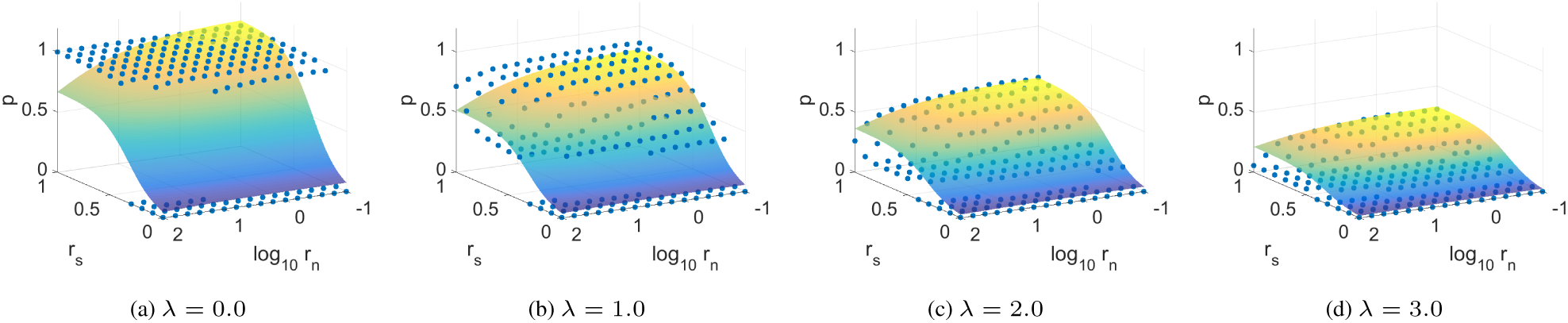
The partial simulation results (points) and fitted surfaces of the proposed model using the spectrum-based open search. In total, 2816 (= 11*×*16*×*16) parameter combinations are simulated. In this figure, 704 (= 11*×*16*×*4) data points are shown. The full simulation results are shown in the Supplementary Notes. When *λ* = 0.0, as shown in (a), there is a clear plateau when *r*_*s*_ is not extremely small. Since no mass shift exists, the spectrum-based open search provides satisfactory results, even though the MS1 tolerance is enlarged to include more modified candidates. As *λ* increases, the plateau gradually drops. When *λ* = 3.0, the maximum probability of obtaining correct identifications is less than 50%.

**Fig. 2:**
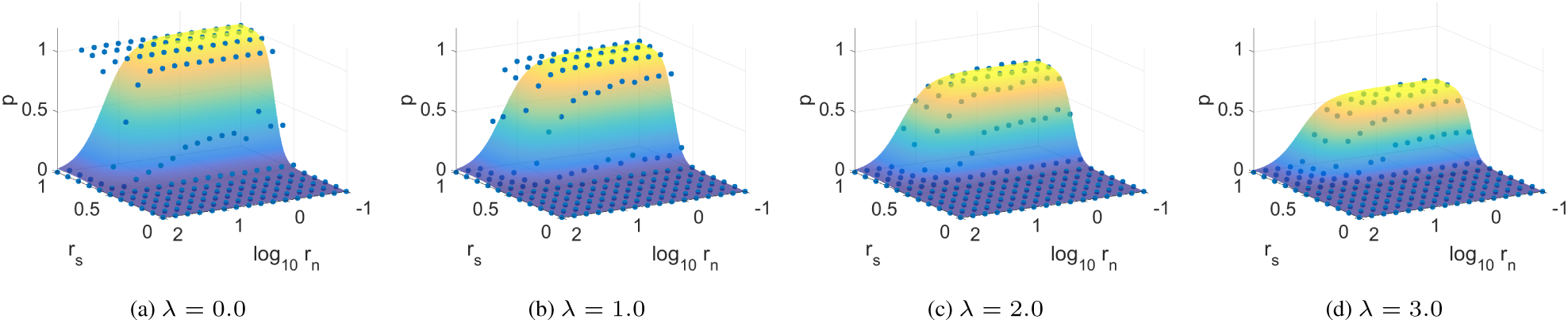
The partial simulation results (points) and fitted surfaces of the proposed model using the tag-based open search. In total, 2816 (= 11 *×* 16 *×* 16) parameter combinations are simulated. In this figure, 704 (= 11*×*16*×*4) data points are shown. The full simulation results are shown in the Supplementary Notes. When *λ* = 0.0, as shown in (a), there is a clear plateau when *r*_*s*_ is greater than 0.6 and *r*_*n*_ is not extremely large. In this region, the tag-based open search will correctly identify unmodified peptides with higher than 90% probability. As *r*_*s*_ decreases and *r*_*n*_ increases, the probability of obtaining correct identifications gradually drops to zero. Also, as *λ* increases, the maximum probability decreases. Compared with Fig. 1, when *λ* is large, the maximum probability of the tag-based open search is greater than that of the spectrum-based open search. This means that the unmodified information captured by the tag vectors can provide more useful information during the identification. However, when *λ* is small, the tag-based open search shows worse robustness than the spectrum-based open search when *r*_*n*_ is high and *r*_*s*_ is small.

As shown in the figures, the probability of obtaining correct identifications achieves the maximum when *r*_*s*_ is largest and *r*_*n*_ is smallest. This meets our expectation because more information about the true peaks and less interference from the noise peaks should benefit the identification process. As *r*_*s*_ decreases and *r*_*n*_ increases, the probability decreases, which means that the performance of open search methods degrades. When *λ* increases, the probability decreases because the existence of PTMs always destroy some patterns of the backbone peptides. Therefore, even though open search methods are designed to tolerate the mass shifts introduced by PTMs, the performance of large *λ* is not as good as the performance of small *λ*.

### 3.2 Model of performance trend

We use a unified model to describe the performance trend of the spectrum-based and tag-based open search methods. The model reads

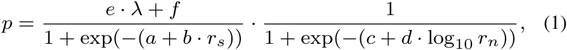

where *p* is the probability of obtaining correct identifications, and *a, b, c, d, e* and *f* are coefficients. The values of *a, b, c, d, e* and *f* will be obtained through model fitting on the simulation results.

To obtain a stable fitted model using Eq. (1), we use a two-step procedure. We compute the values of *e* and *f* using least squares first. Then, we compute the values of *a, b, c* and *d* using nonlinear least squares with the fixed *e* and *f*. The details of the fitting procedure are shown in the Supplementary Notes. The fitted models are shown as separate surfaces in Fig. 1 and Fig. 2. The fitted coefficients are shown in Table 1.

**Table 1.** Fitted coefficients of the model of performance trend (Eq. (1)).

| Coefficient | $a$ | $b$ | $c$ | $d$ | $e$ | $f$ |
| --- | --- | --- | --- | --- | --- | --- |
| Spectrum-based | -2.6026 | 7.9758 | 3.1748 | -1.3178 | -0.2420 | 1.0603 |
| Tag-based | -16.2532 | 23.9516 | 5.9019 | -4.7211 | -0.1516 | 1.0149 |

### 3.3 Evaluation of the model

We check the sum-of-squares error (SSE) of the fitted model. The SSE of the model for the spectrum-based open search is 62.3037, and the corresponding root-mean-square error (RMSE) is 0.14874. The SSE of the model for the tag-based open search is 24.4097, and the corresponding RMSE is 0.093103. The absolute residual histograms of the fitted model for both types of open search are shown in Fig. 3 and Fig. 4. From the histograms, we find that the majority of the data points are fitted well.

**Fig. 3:**
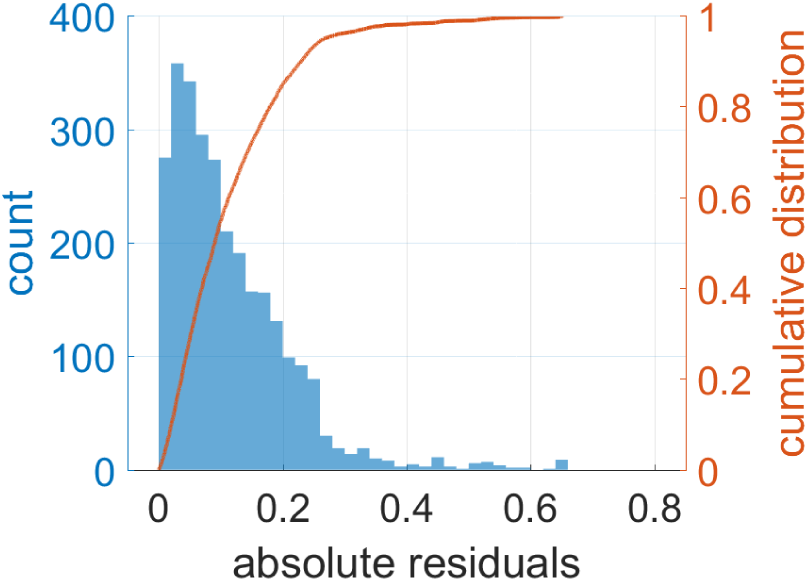
The histogram of the absolute residuals of the fitted model for the spectrum-based open search. The histogram is plotted using a bin width of 0.02. Compared with the tag-based fitting result, the spectrum-based fitting has a larger error.

**Fig. 4:**
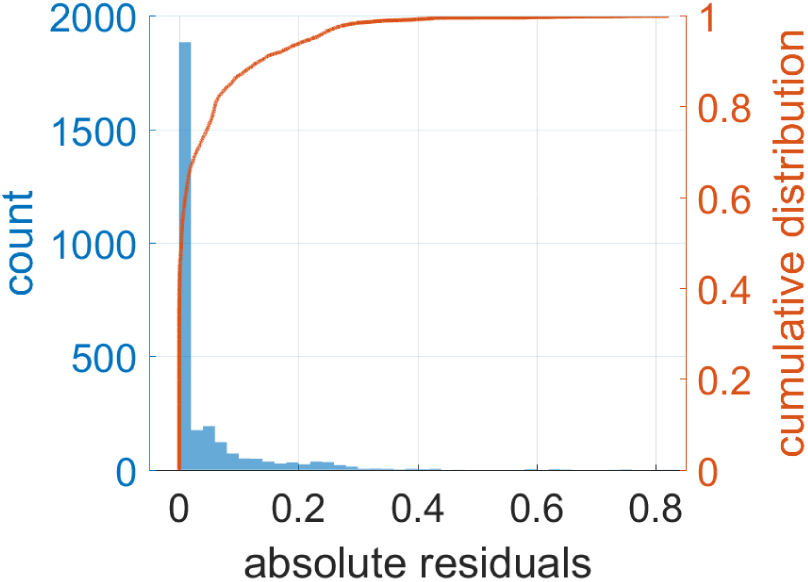
The histogram of the absolute residuals of the fitted model for the tag-based open search. The histogram is plotted using a bin width of 0.02. The majority of the residuals are within *±*0.02 (roughly 60% in the first bin). From manual examination of large residuals, we find that most of them come from the transition from small *r*_*n*_ to large *r*_*n*_ when *r*_*s*_ is large (see Fig. 2).

To evaluate the analytical model, we generate an independent testing dataset. We randomly choose 400 parameter combinations in the same parameter space used above (i.e., uniformly sampling random (*r*_*s*_, *r*_*n*_, *λ*) subject to *r*_*s*_ ∈ [0, 1], *r*_*n*_ ∈ [0.1, 100], *λ ∈* [0.0, 3.75]). We also examine the absolute residuals by predicting the probability values using the fitted model. Histograms are shown in Fig. 5 and Fig. 6. They are similar to Fig. 3 and Fig. 4.

**Fig. 5:**
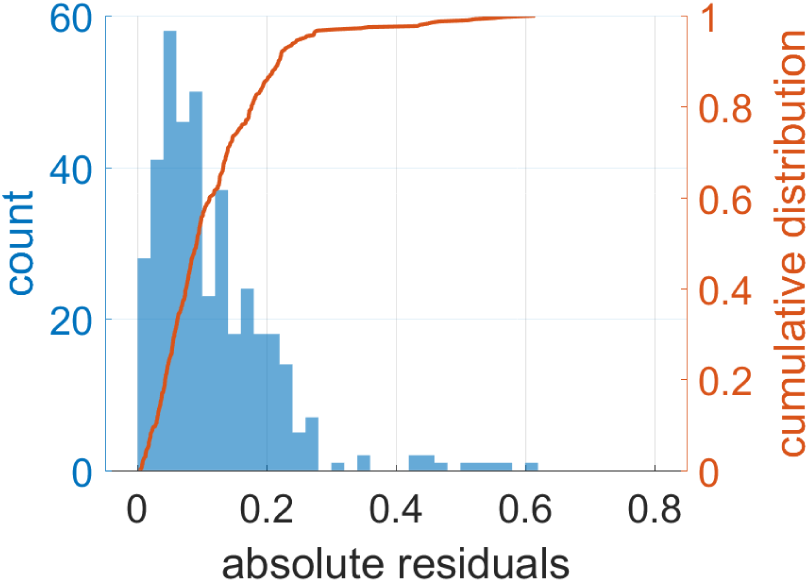
The histogram of the absolute residuals of the model for the spectrum-based open search using an independent testing dataset. The histogram is plotted using a bin width of 0.02. In total, 400 parameter combinations are sampled. This histogram is similar to Fig. 3.

**Fig. 6:**
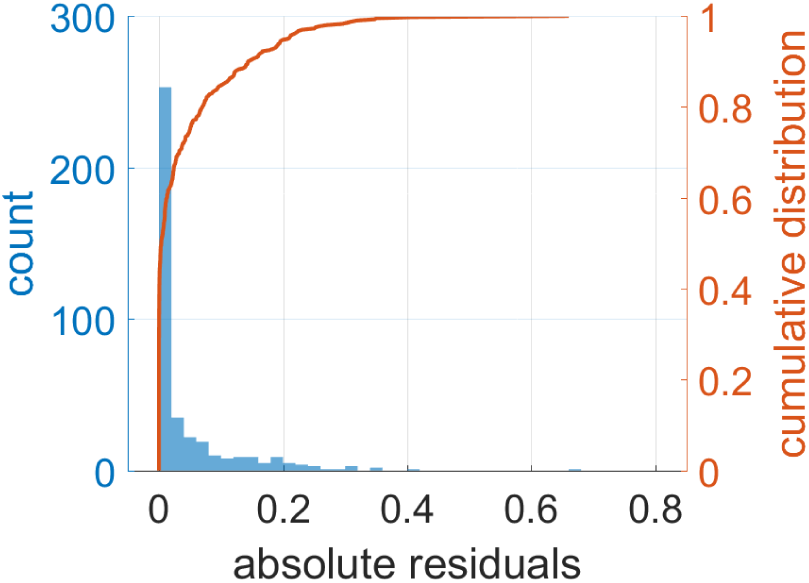
The histogram of the absolute residuals of the model for the tag-based open search using an independent testing dataset. The histogram is plotted using a bin width of 0.02. In total, 400 parameter combinations are sampled. This histogram is similar to Fig. 4.

We further generate a testing dataset with higher noise levels to test the extrapolation performance of our model. Because increasing the noise level drastically increases the computing time in the simulations, we only choose 10 parameter combinations in this dataset. These parameter combinations are randomly drawn from *r*_*s*_ = 0 to *r*_*s*_ = 1 (the whole range of *r*_*s*_), *r*_*n*_ = 100 to *r*_*n*_ = 316.23 (higher noise levels), and *λ* = 0.0 to *λ* = 3.75. We list these parameter combinations in Table 2 together with the empirical and predicted probabilities of obtaining correct identifications. The corresponding absolute residual histograms are shown in Fig. 7 and Fig. 8. Although we have not included any parameter combinations with *r*_*n*_ *>* 10^2^ for model fitting, the fitted model can correctly extrapolate the performance trend.

**Table 2.**
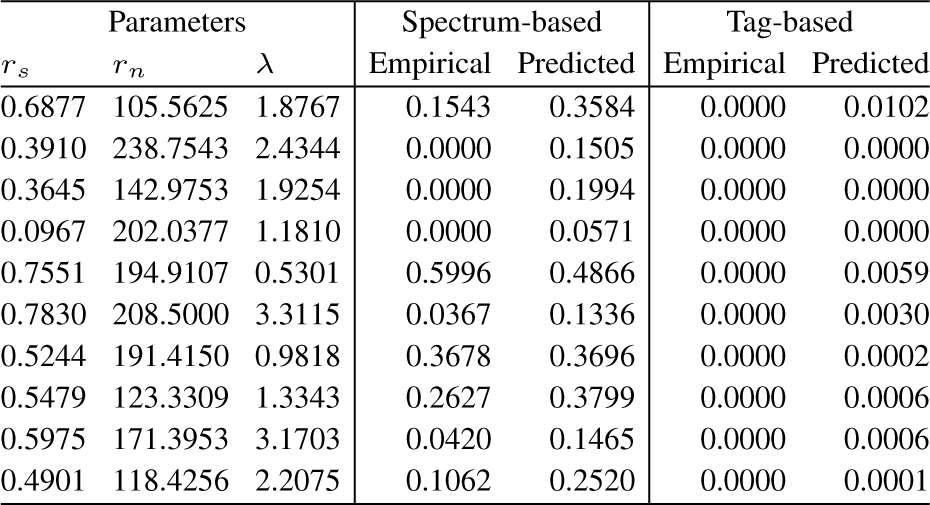
The empirical and predicted probabilities of obtaining correct identifications in a testing dataset with higher noise levels.

| Parameters |  |  | Spectrum-based |  | Tag-based |  |
| --- | --- | --- | --- | --- | --- | --- |
| $r_s$ | $r_n$ | $\lambda$ | Empirical | Predicted | Empirical | Predicted |
| 0.6877 | 105.5625 | 1.8767 | 0.1543 | 0.3584 | 0.0000 | 0.0102 |
| 0.3910 | 238.7543 | 2.4344 | 0.0000 | 0.1505 | 0.0000 | 0.0000 |
| 0.3645 | 142.9753 | 1.9254 | 0.0000 | 0.1994 | 0.0000 | 0.0000 |
| 0.0967 | 202.0377 | 1.1810 | 0.0000 | 0.0571 | 0.0000 | 0.0000 |
| 0.7551 | 194.9107 | 0.5301 | 0.5996 | 0.4866 | 0.0000 | 0.0059 |
| 0.7830 | 208.5000 | 3.3115 | 0.0367 | 0.1336 | 0.0000 | 0.0030 |
| 0.5244 | 191.4150 | 0.9818 | 0.3678 | 0.3696 | 0.0000 | 0.0002 |
| 0.5479 | 123.3309 | 1.3343 | 0.2627 | 0.3799 | 0.0000 | 0.0006 |
| 0.5975 | 171.3953 | 3.1703 | 0.0420 | 0.1465 | 0.0000 | 0.0006 |
| 0.4901 | 118.4256 | 2.2075 | 0.1062 | 0.2520 | 0.0000 | 0.0001 |

**Fig. 7:**
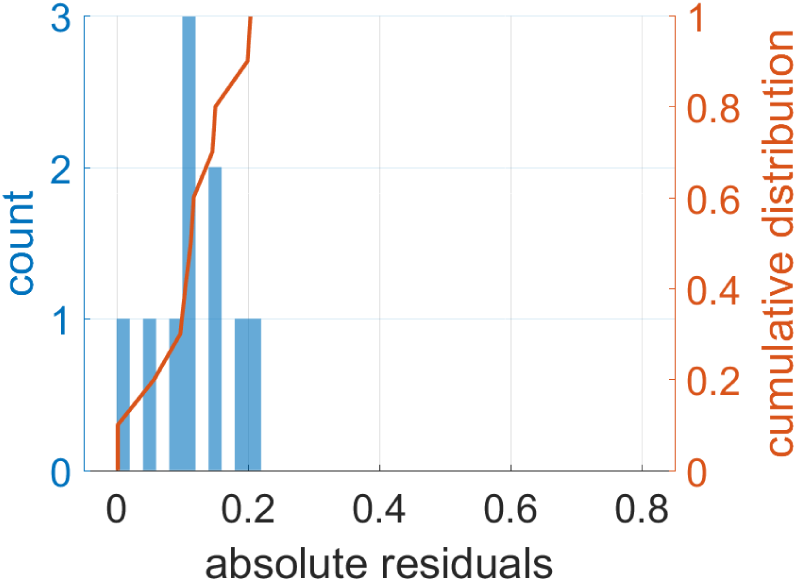
The histogram of the absolute residuals of the model for the spectrum-based open search using a testing dataset with higher noise levels. The histogram is plotted using a bin width of 0.02. In total, 10 parameter combinations are sampled.

**Fig. 8:**
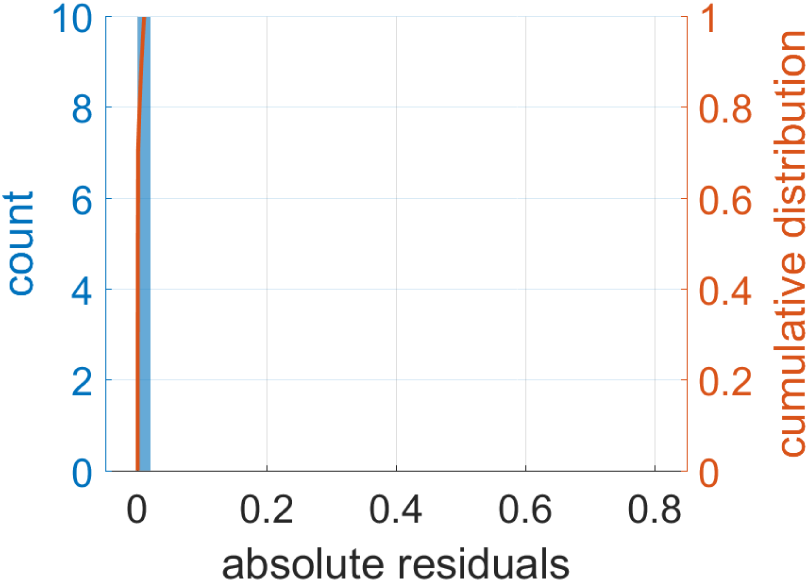
The histogram of the absolute residuals of the model for the tag-based open search using a testing dataset with higher noise levels. The histogram is plotted using a bin width of 0.02. In total, 10 parameter combinations are sampled. It provides good predictions under the condition when the noise level is high. In this area, the tag-based open search usually shows a probability close to zero.

The absolute residual histograms of the spectrum-based open search (Fig. 3, Fig. 5 and Fig. 7) show a similar pattern, though the sample size in Fig. 7 is relatively small. Compared with the absolute residual histograms of the tag-based open search (Fig. 4, Fig. 6 and Fig. 8), the histograms of the spectrum-based open search have larger errors. By viewing the fitted surfaces in Fig. 1, we find that when *r*_*s*_ decreases from a large value, there is a steep transition. As a result, the surfaces do not fit well. From Table 2, we find that the 7th row shows a good match between the empirical probability and the predicted probability for the spectrum-based open search, while the others show larger errors. Also, similar to the histograms in Fig. 3 and Fig. 5, most of the absolute residuals in Table 2 are within *±*0.2.

The absolute residual histograms of the tag-based open search (Fig. 4, Fig. 6 and Fig. 8) show a better match. Roughly 60% of the absolute residuals are in the first bin (i.e., within *±*0.02), and roughly 80% of the absolute residuals are within *±*0.1. The reason of the better match of the tag-based open search is that the simulation results of the tag-based open search show a clearer area with high probabilities. As a result, the proposed model with two sigmoid functions can describe this scenario better.

## 4 Discussion

In this paper, we systematically studied the performance trend of open search methods with respect to the number of PTMs in peptides, the ratio of missing peaks and the noise level. Based on the analytical model we fitted using a simulation dataset, we can reveal the necessary condition of open search methods. For example, if we consider *p ≥* 70% as an acceptable threshold of performance, the corresponding feasible parameter range can be obtained from

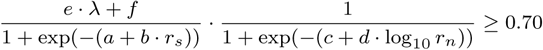

subject to an implicit constraint 0 *≤ r*_*s*_ *≤* 1. When we use the tag-based open search, and would like to identify peptides with at most 2 PTMs, the feasible region will be the shaded area in Fig. 9. Under this setting, the condition of *r*_*s*_ = 0.9 and *r*_*n*_ = 1.0 is within this feasible region, but the condition of *r*_*s*_ = 0.85 and *r*_*n*_ = 10.0 is outside this feasible region.

**Fig. 9:**
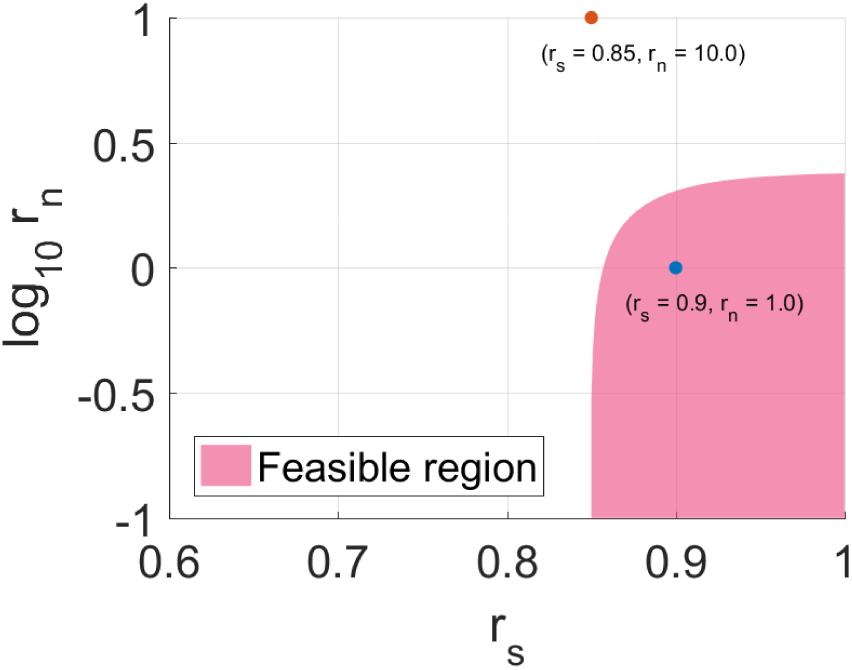
An example of the necessary condition of the tag-based open search when specifying 70% as an acceptable threshold in the identification of peptides with at most 2 PTMs.

When *λ* is small, i.e., the expected number of PTMs in the spectra is small, as shown in Fig. 1 and Fig. 2, the spectrum-based open search performs more robustly than the tag-based open search because it tolerates more missing peaks in a spectrum. However, when *λ* is large, the tag-based open search is better, because it can achieve a higher maximum probability of obtaining correct identifications when *r*_*s*_ is high and *r*_*n*_ is small. If *r*_*s*_ and *r*_*n*_ can be well controlled in the experiment, the tag-based open search would be preferred because it shows a better performance compared with the spectrum-based open search, especially when *λ* is large. When the experiment conditions are not well controlled, the tag-based open search may not be robust enough to accommodate different scenarios.

In our simulation, intensity information in the experimental spectra and dissociation patterns around the matched ions are not fully utilized. In addition, the uniform assumption of noise peak positions may not match the real cases, as some noise peaks may come from certain unknown molecules. Therefore, the analytical model tends to underestimate the effect of a specified noise level. The improvement of scoring functions may pull the surfaces up, but the overall trend in the proposed analytical model should stay similar.

