## Supplementary Materials for "Understanding the limit of open search in the identification of peptides with post-translational modifications — A simulation-based study"

March 27, 2018

#### Contents

**1 Ideal Case Examination** **2**

**2 Full Simulation Results** **3**

**3 Model Fitting Procedure** **6**

*Jiaan Dai and Fengchao Yu contributed equally to this work.

### 1 Ideal Case Examination

To analyze the intrinsic limit of the spectrum vectors and the tag vectors in open search methods, we conduct simulations under the ideal setting. We use the perfect spectrum vector and tag vector given a base peptide for the simulated spectrum, which are the theoretical spectrum of the base peptide and the theoretical tag set of the base peptide, respectively. Instead of uniformly choosing base peptides, we enumerate every peptides in the database to provide a thorough analysis. We run simulations on the following 3 databases: (1) the *E.coli* K12 protein database (downloaded on 2016.10.19 from UniProtKB/SwissProt, 4434 proteins), (2) the *S. cerevisiae* protein database (downloaded on 2016.10.19 from UniProtKB/SwissProt, 6721 proteins), and (3) the *Homo sapiens* (human) protein database (downloaded on 2016.03.10 from UniProtKB/SwissProt, 20198 proteins). The databases are digested with the trypsin cleavage rule, and no missed cleavages are allowed. The allowed range of the peptide mass is $[700, 5000]$ Da, and the MS1 tolerance is 250 Da. We don't conduct FDR control in this analysis, and we count an identification as correct if its label can be selected by the highest similarity rule unambiguously. The numbers of correct identifications are shown in Table 1 and Table 2.

Table 1: The number of correct identifications in ideal case simulations using the spectrum-based open search. The percentage is calculated by #Correct / #Spectrum.

| Database | #Spectrum | #Correct | Percentage |
| --- | ---: | ---: | ---: |
| *E. coli* | 70105 | 70105 | 100% |
| *S. cerevisiae* | 154006 | 154006 | 100% |
| *Homo sapiens* | 554376 | 554376 | 100% |

Table 2: The number of correct identifications in ideal case simulations using the tag-based open search. The percentage is calculated by #Correct / #Spectrum.

| Database | #Spectrum | #Correct | Percentage |
| --- | ---: | ---: | ---: |
| *E. coli* | 70105 | 70103 | 99.997% |
| *S. cerevisiae* | 154006 | 153909 | 99.937% |
| *Homo sapiens* | 554376 | 553783 | 99.893% |

The percentages of correct identifications are close to 100%, which indicates that the representations of both the spectrum vector and the tag vector will not significantly affect the capability of assigning correct labels when perfect spectra are available, though the representation of tag vector may fail to distinguish two peptides in some rare cases.

### 2 Full Simulation Results

The full simulation results are shown in Figure 1 and Figure 2. The probability of obtaining correct identifications achieves the maximum when $r_s$ is largest and $r_n$ is smallest. This meets our expectation because more information about the true peaks and less interference from the noise peaks should benefit the identification process. As $r_s$ decreases and $r_n$ increases, the probability decreases, which means that the performance of open search methods degrades. When $\lambda$ increases, the probability decreases because the existence of PTMs always destroys some patterns of the backbone peptides. Therefore, even though open search methods are designed to tolerate the mass shifts introduced by PTMs, their performance with large $\lambda$ is not as good as their performance with small $\lambda$.

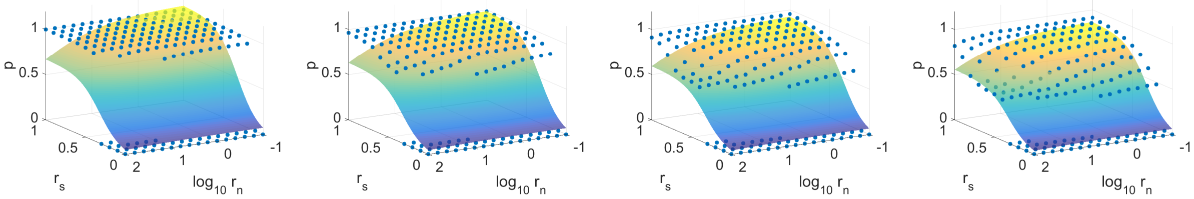

(a) $\lambda = 0.0$ (b) $\lambda = 0.25$ (c) $\lambda = 0.5$ (d) $\lambda = 0.75$

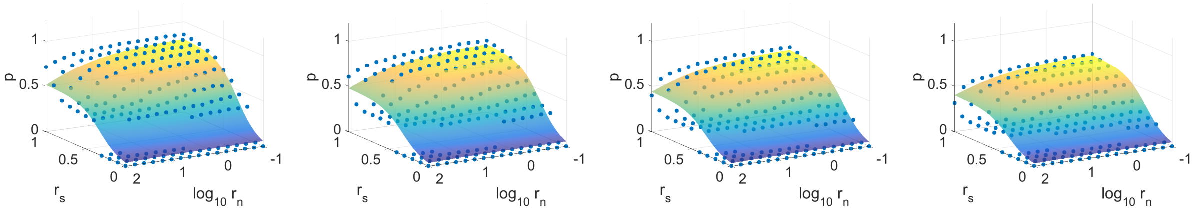

(e) $\lambda = 1.0$ (f) $\lambda = 1.25$ (g) $\lambda = 1.5$ (h) $\lambda = 1.75$

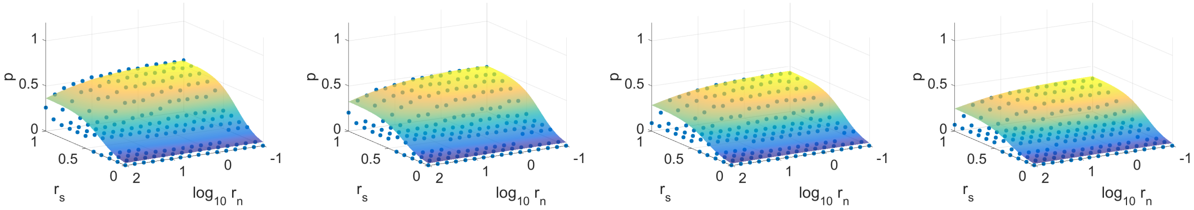

(i) $\lambda = 2.0$ (j) $\lambda = 2.25$ (k) $\lambda = 2.5$ (l) $\lambda = 2.75$

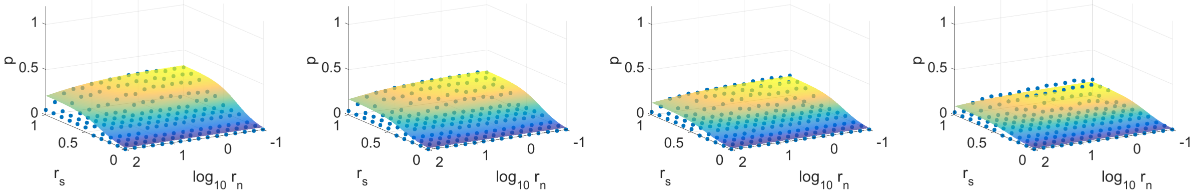

(m) $\lambda = 3.0$ (n) $\lambda = 3.25$ (o) $\lambda = 3.5$ (p) $\lambda = 3.75$

Figure 1: The simulation results (points) and fitted surfaces of the proposed model using spectrum-based open search. In total, 2816 ($= 11 \times 16 \times 16$) parameter combinations are simulated. When $\lambda = 0.0$, as shown in (a), there is a clear plateau when $r_s$ is not extremely small. Since no mass shift exists, the spectrum-based open search provides satisfactory results even though the MS1 tolerance is enlarged to include more modified candidates. As $\lambda$ increases, the plateau gradually drops. When $\lambda$ is greater than or equal to 2.5, the maximum probability it can achieve in the simulated space of $r_s$ and $r_n$ is less than 50%.

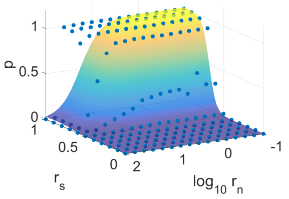


(a) $\lambda = 0.0$ (b) $\lambda = 0.25$ (c) $\lambda = 0.5$ (d) $\lambda = 0.75$

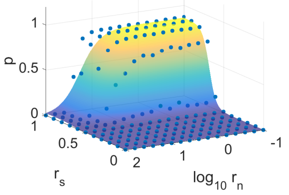


(e) $\lambda = 1.0$ (f) $\lambda = 1.25$ (g) $\lambda = 1.5$ (h) $\lambda = 1.75$

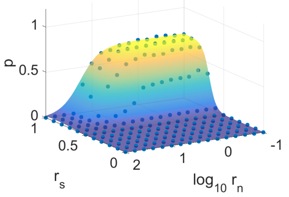


(i) $\lambda = 2.0$ (j) $\lambda = 2.25$ (k) $\lambda = 2.5$ (l) $\lambda = 2.75$

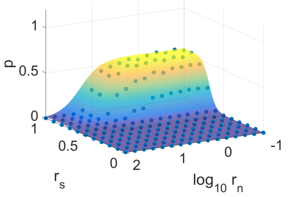


(m) $\lambda = 3.0$ (n) $\lambda = 3.25$ (o) $\lambda = 3.5$ (p) $\lambda = 3.75$

Figure 2: The simulation results (points) and fitted surfaces of the proposed model using tag-based open search. In total, 2816 ($= 11 \times 16 \times 16$) parameter combinations are simulated. When $\lambda = 0.0$, as shown in (a), there is a clear plateau when $r_s$ is greater than 0.6 and $r_n$ is not extremely large. In this region, the tag-based open search will correctly identify unmodified peptides with higher than 90% probability. As $r_s$ decreases and $r_n$ increases, the probability of obtaining correct identifications gradually drops to zero. Also, as $\lambda$ increases, the maximum probability in the plateau decreases. Compared with Figure 1, when $\lambda$ is large, the maximum probability shown in the results of the tag-based open search is greater than that shown in the results of the spectrum-based open search, which means that the unmodified information captured by the tag vectors can provide more useful information during the identification. However, when $\lambda$ is small, the tag-based open search shows worse robustness when $r_n$ is high and $r_s$ is small.

### 3 Model Fitting Procedure

To fit the analytical model, we first use a linear function

$$e \cdot \lambda + f = p_{\max}^{(\lambda)} \tag{1}$$

to describe the relationship between the maximum probability and the given $\lambda$ condition. To obtain $p_{\max}^{(\lambda)}$, we collect the maximum probabilities in the simulation results by fixing the value of $\lambda$. Therefore, we are able to obtain the raw trend of $p_{\max}^{(\lambda)}$ versus $\lambda$, and we compute the coefficients $e$ and $f$ in Eq (1) using the least squares estimation. Coefficients $e$ and $f$ are fixed when we compute the values of other coefficients. After plugging $e$ and $f$ into the model, we use the nonlinear least squares solver (with the Trust-Region algorithm) in the Curve Fitting Toolbox in MATLAB to obtain the values of $a, b, c$ and $d$. The trend of $p_{\max}^{(\lambda)}$ versus $\lambda$ of the spectrum-based open search is shown in Figure 3, and the trend of $p_{\max}^{(\lambda)}$ versus $\lambda$ of the tag-based open search is shown in Figure 4, for validating the linearity of Eq (1). The linearity is clear between $p_{\max}^{(\lambda)}$ and $\lambda$ when $\lambda \leq 3.75$.

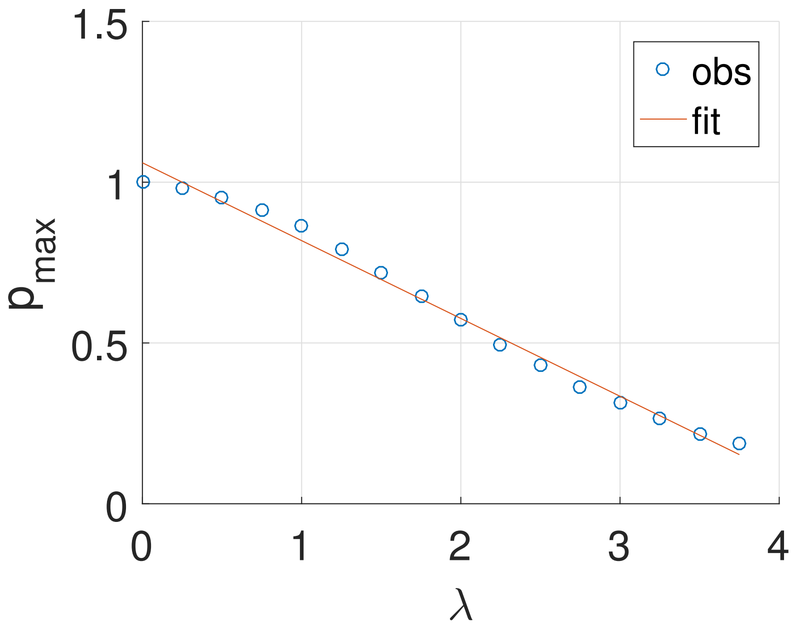


Figure 3: $p_{\max}^{(\lambda)}$ versus $\lambda$ when $\lambda \leq 3.75$ using the spectrum-based open search. The graph shows a good linear relationship between the empirical maximum probability and the expected number of random mass shifts in the simulation. After conducting least squares on $p_{\max}^{(\lambda)} = e \cdot \lambda + f$, we fix the values of $e$ and $f$ in the model, and conduct nonlinear least squares to find the values of $a, b, c$ and $d$.

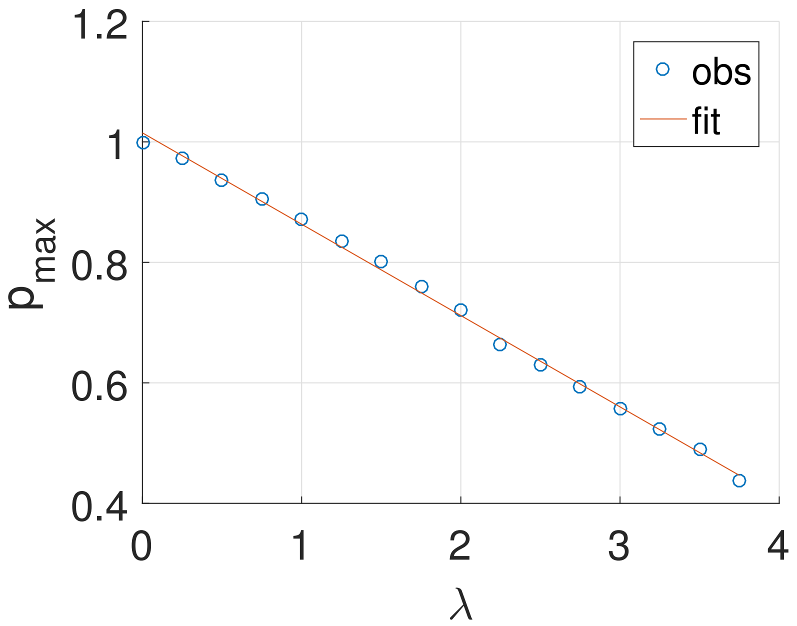


Figure 4: $p_{\max}^{(\lambda)}$ versus $\lambda$ when $\lambda \leq 3.75$ using the tag-based open search. The graph shows a clear linear relationship between the empirical maximum probability and the expected number of random mass shifts in the simulation. After conducting least squares on $p_{\max}^{(\lambda)} = e \cdot \lambda + f$, we fix the values of $e$ and $f$ in the model, and conduct nonlinear least squares to find the values of $a, b, c$ and $d$.
